# Reinforcement learning during locomotion

**DOI:** 10.1101/2023.09.13.557581

**Authors:** Jonathan M Wood, Hyosub E Kim, Susanne M Morton

**Affiliations:** Department of Physical Therapy, University of Delaware, Newark, DE 19713, United States; Interdisciplinary Graduate Program in Biomechanics & Movement Science, University of Delaware, Newark, DE 19713, United States; Department of Psychological and Brain Sciences, University of Delaware, Newark, DE 19716, United States; School of Kinesiology, University of British Columbia, Vancouver, BC, Canada

## Abstract

When learning a new motor skill, people often must use trial and error to discover which movement is best. In the reinforcement learning framework, this concept is known as exploration and has been linked to increased movement variability in motor tasks. For locomotor tasks, however, increased variability decreases upright stability. As such, exploration during gait may jeopardize balance and safety, making reinforcement learning less effective. Therefore, we set out to determine if humans could acquire and retain a novel locomotor pattern using reinforcement learning alone. Young healthy male and female participants walked on a treadmill and were provided with binary reward feedback (indicated by a green checkmark on the screen) that was tied to a fixed monetary bonus, to learn a novel stepping pattern. We also recruited a comparison group who walked with the same novel stepping pattern but did so by correcting for target error, induced by providing real time veridical visual feedback of steps and a target. In two experiments, we compared learning, motor variability, and two forms of motor memories between the groups. We found that individuals in the binary reward group did, in fact, acquire the new walking pattern by exploring (increasing motor variability). Additionally, while reinforcement learning did not increase implicit motor memories, it resulted in more accurate explicit motor memories compared to the target error group. Overall, these results demonstrate that humans can acquire new walking patterns with reinforcement learning and retain much of the learning over 24 hours.

**Significance Statement:** Humans can learn some novel movements by independently discovering the actions that lead to success. This discovery process, exploration, requires increased motor variability to determine the best movement. However, in bipedal locomotion especially, increasing motor variability decreases stability, heightening the risk of negative outcomes such as a trip, injury, or fall. Despite this stability constraint, the current study shows that individuals do use exploration to find the most rewarding walking patterns. This form of learning led to improved explicit retention but not implicit aftereffects. Thus, the reinforcement learning framework can explain findings across a wide range of motor and cognitive tasks, including locomotion.

## Introduction

In the 2020 Summer Olympics, U.S. athlete Sydney McLauchlin broke the world record and won the gold medal in the 400-meter hurdles, aided by her proficiency in jumping over the hurdles equally well with both her dominant and non-dominant legs. While reward (clearing the hurdle) is ostensibly a critical learning signal for this and other tasks, its specific role in locomotor learning remains unknown. Rewards influence learning via reward prediction errors, the difference between expected and actual rewards (Rescorla and Wagner, 1972). These error signals form the basis for reinforcement learning theory, a widely used framework describing how animals and machines learn the values of actions (Sutton and Barto, 2017). A core component of reinforcement learning is exploration, defined as sampling from a range of possible actions to update their values. In movement-based reinforcement learning, one sign of exploration is increased motor variability above baseline motor noise as this likely aids in motor learning (Cashaback et al., 2019, 2017; Dhawale et al., 2017; Therrien et al., 2018, 2016). Thus, we refer to exploration in the context of motor learning as an increase in motor variability above baseline motor noise. However, in locomotion, the cost of changing motor variability is much higher, as too much or too little variability in gait patterns reduces stability and could lead to catastrophic consequences like loss of balance, falls, or injuries (Brach et al., 2005; Hausdorff et al., 2001, 1997; Maki, 1997; McAndrew Young and Dingwell, 2012) .

During motor tasks, reinforcement learning is often probed by providing only binary reward feedback, and has been demonstrated in reaching (Izawa and Shadmehr, 2011), speech production (Parrell, 2021), and saccadic eye movements (Madelain et al., 2011). Since binary reward feedback does not provide information about the direction or distance from a task goal, the only signal by which to learn the correct movement is a reward prediction error, resulting in exploratory behavior. In sharp contrast, learning signals such as target error (i.e., the difference between a task goal and movement feedback) and sensory prediction error (i.e., the difference between the expected and actual sensory consequences of movements), convey both magnitude and directional information. As a result, they produce comparatively less exploration than reward prediction errors (Cashaback et al., 2019; Izawa and Shadmehr, 2011; Therrien et al., 2016). To date, no studies have determined if binary reward feedback can produce learning, and the expected exploratory behavior, during locomotion.

Reinforcement learning also impacts motor memories. Implicit aftereffects, defined as motor memories not under conscious control (Krakauer et al., 2019), are strengthened when reinforcement is combined with sensorimotor adaptation (Galea et al., 2015; Huang et al., 2011; Shmuelof et al., 2012), or use-dependent learning (Mawase et al., 2017; Bao and Lei, 2022; c.f., Tsay et al., 2022). Reinforcement learning may also strengthen explicit retention, defined as the ability to consciously remember and reproduce a previously learned movement (Schmidt and Lee, 2005), potentially because individuals benefit from determining what the successful movement is themselves through exploration, improving engagement in the task, unlike when receiving full visual feedback of movements (Hasson et al., 2015; Winstein et al., 1994; Winstein and Schmidt, 1990). However, previous studies examining explicit retention in reinforcement learning either did not induce learning with purely binary reward feedback (Codol et al., 2018; Hasson et al., 2015), or did not include a comparison group (Holland et al., 2018). Therrien et al. (2016) found retention was better after binary reward feedback than sensory prediction and target errors, but this probe into retention may have also included a strong implicit component. Therefore, the extent to which reinforcement learning alone contributes to implicit and explicit motor memory formation remains unclear.

The primary objective of this study was to determine if individuals can acquire a novel gait pattern using reinforcement learning. We tested this by providing binary reward feedback to participants learning to walk with a longer left step length than baseline. For comparison, we recruited a group that performed the same walking pattern using explicit, target error feedback, which does not induce significant exploration. Prior work has demonstrated that use-dependent learning is the predominant source of implicit aftereffects in this target error paradigm (Wood et al., 2021, 2020), and neither implicit aftereffects nor explicit retention is contaminated by sensory prediction errors. We hypothesized that the reward prediction error group would learn the new walking pattern using greater exploration (a greater increase in motor variability above baseline), and that they would have larger implicit aftereffects and more accurate explicit retention compared to the target error group.

## Materials & Methods

### Participants

A total of 59 (34 female, 25 male) young, healthy adults were recruited from the University of Delaware community. Within 2 separate experiments, participants were randomized into 1 of 2 groups: a reward prediction error (RPE) group and a target error (TE) group. Participants were excluded if they had any orthopedic or neurologic conditions that might interfere with their ability to learn a new walking pattern. Five participants were removed from the analysis (2 due to technical errors, 2 due to not following instructions, and 1 who was provided an erroneous verbal reminder of the instructions during learning). Thus, thirty participants were included in experiment 1 and 24 in experiment 2. Before participating, all individuals signed a written, informed consent document. This study was approved by the University of Delaware Institutional Review Board.

### Experimental design

#### Feedback

Participants walked on a dual belt treadmill (Figure 1A; Bertec, Columbus OH) at a self-selected, comfortable but brisk pace, constrained between 1.0 and 1.2 m/s (Wood et al., 2021, 2020). This allowed each participant to walk at a comfortable pace given their biomechanical constraints, preventing differences in task difficulty due to treadmill speed. The belts moved at the same speed throughout the experiment. Participants wore a ceiling mounted harness, which did not provide body weight support, and lightly held onto a handrail for safety while walking. A computer monitor placed in front of the treadmill provided real-time feedback of participants’ left step lengths (The Motion Monitor Toolbox, Innovative Sports Training Inc., Chicago IL). We chose to focus on step length because it is a robustly modifiable gait parameter that has frequently been the focus of locomotor learning paradigms (French et al., 2021, 2018; Reisman et al., 2005; Roemmich et al., 2016; Wood et al., 2021, 2020), and because both implicit and explicit forms of motor memory have been observed after learning an altered step length pattern (French et al., 2021, 2018; Wood et al., 2021, 2020).

**Figure 1:**
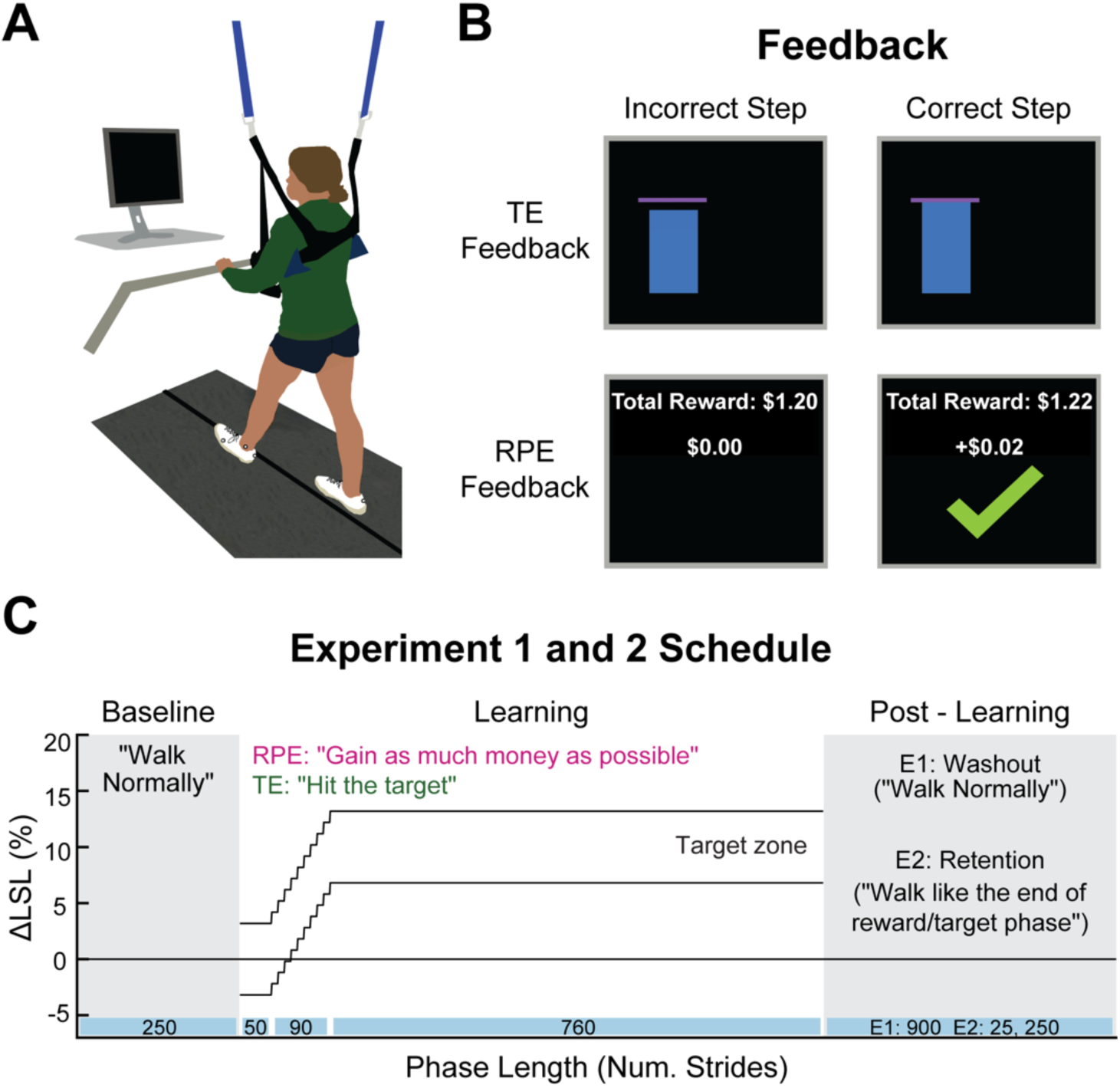
Experimental paradigm. **(A)** All participants walked on a dual-belt treadmill, at a comfortable self-selected pace with a computer monitor in front of them. **(B)** The RPE and TE groups received different feedback during the learning phase. The RPE group received only binary reward feedback, with a check mark and money added to a total when they performed a correct step length. The TE group received real-time feedback of their left step length related to the pink, horizontal target line. **(C)** All participants walked in 3 different phases: 1) baseline, where individuals were asked to walk normally; 2) learning, where they gradually learned to walk with a longer left step length with the feedback and instructions depending on group assignment; 3) post-learning, where implicit aftereffects (experiment 1) or explicit retention (experiment 2) were probed, both without visual feedback. Participants in experiment 2 also were tested for explicit retention 24 hours later (not shown). Note that the target window in this figure was taken from a representative participant and is in terms of ΔLSL, but for all participants the target window was ±2 cm.

We compared a group that received reward prediction error feedback (RPE group) to a group that received explicit task instructions and visual target errors (TE group). The TE group received real-time visual feedback of their step length relative to a step length target (Figure 1B, top row). The left step length appeared as a blue bar, starting with a height of ‘0’ and growing vertically during the swing phase, then holding position on the screen at its final height at the time of heel strike, until the next swing phase began. The TE group was instructed in the definition of a step length and that they needed to hit a pink horizontal target line exactly with the blue bar that represented their left step length. They were also informed how the target left step length would change during learning. Although they could see if they hit the target line, no external reward was provided either on the screen or verbally by the experimenter when they successfully hit the target. Thus, since the TE group knew exactly where to step and could correct for any target errors throughout the task, they had no need to explore. Furthermore, since the feedback was veridical, sensory prediction errors could not contaminate either implicit aftereffects or explicit retention.

The RPE group was provided with binary reward feedback (Figure 1B, bottom row) that was tied to a fixed monetary bonus. This feedback took the form of a large green check mark and money added to a total when a left step length fell within the target window, the same target as the TE group, but here, invisible to participants. If successful, the check mark appeared on the screen at heel strike and remained on the screen until the next swing phase began. When the left step length fell outside of the target window, no check mark appeared, and no money was added to the total. A running tally of the total reward earned was consistently displayed at the top of the screen during the learning phase. The RPE group was told that their goal was to gain as much money as possible. As awareness of the rewarded parameter may be important for movement based reinforcement learning (Manley et al., 2014), individuals in the RPE group were told that the reward was contingent on their left step length. The experimenter then explained and demonstrated a left step length until the participant could explain the concept themselves.

Based on pilot testing, we also told the participants in the RPE group to start off by walking normally so they could achieve some success at the beginning of the learning phase, but critically, no information was provided regarding if or how the rewarded step length would change. Thus, to learn the longer left step length, the RPE group was provided with no other learning signal except a reward prediction error, the difference between the expected and actual reward. Participants in both groups were provided with a brief, verbal reminder of their instructions every 100 strides (TE: “hit the target”; RPE: “gain as much money as possible”).

Participants in the RPE group were told that they would earn a base dollar amount ($10) per visit, plus they had the potential to earn additional “bonus” money, based upon how well they performed the task (maximum $18 bonus). However, in actuality, all participants earned the same amount, which was the base amount plus the maximum bonus amount, regardless of task performance. Participants in the TE group were told that they would earn a base dollar amount per visit, which was equivalent to the amount paid to participants in the RPE group. This payment method was chosen, with IRB consultation, to ensure fairness, i.e., to prevent differences in compensation that depended on group assignment or learning abilities.

#### Experimental schedules

Participants in both groups walked under three different phases: baseline, learning and post-learning (Figure 1C). During baseline walking (250 strides) participants were asked to walk normally and no visual feedback was provided. Next, participants performed a learning phase (900 strides), where they gradually learned to lengthen their left step length by 10% from their baseline using the feedback displayed (step length bars and target lines for the TE group or check marks and tally of money earned for the RPE group) on the computer monitor. While the feedback was different between the groups, the learning schedule was the same. For the first 50 strides of the learning phase, the target was centered on each individual’s baseline left step length. The target step length then increased by 1% every 10 strides until it reached its final 10% change from baseline where it remained for the rest of learning (strides 141 to 900). The target maintained a ±2 cm margin of error throughout the learning phase. Thus, while the center of the target was offset by a portion of each participant’s baseline step length, the width of the target was the same for everyone which prevented any differences in difficulty of the task due to baseline step lengths. This target window was well within baseline left step length variability for participants in our sample.

The post-learning phases were used to probe either implicit aftereffects (experiment 1) or explicit retention (experiment 2). Therefore, the visual feedback was turned off during this phase for both experiments, but the instructions were slightly different. In experiment 1, participants completed a washout phase (900 strides) where they were asked to look forward and “walk normally”. Any left step length differences from baseline during this phase represented the implicit aftereffect, a motor memory not under conscious control. In experiment 2, participants completed two explicit retention tests, one immediately after learning (25 strides) and one 24-hours later (250 strides). For both tests, participants were instructed to remember how they walked at the end of the learning phase and repeat that walking pattern exactly. Thus, explicit retention measured the ability to consciously remember and reproduce the previously learned step length pattern, exactly. For both experiments, participants in the RPE group were told that no additional money could be earned during these periods.

### Data collection

Step length was calculated in real time using The Motion Monitor which received kinematic and kinetic data from Nexus software (Vicon Motion Systems Inc., London, UK). Kinetic data, sampled at 1000 Hz, was acquired from the instrumented force plates through each treadmill belt (Bertec, Columbus OH), and kinematic data, sampled at 100 Hz, was acquired from an 8-camera Vicon MX40 motion capture system with Nexus software. To capture the kinematic data, we used a custom marker set with seven retroreflective markers, one on each heel, lateral malleolus, and fifth metatarsal head, and one on the left first metatarsal. The heel markers were used to determine left step length by calculating the difference along the laboratory’s y-axis (anterior-posterior dimension) between the left and the right heel markers at the moment of left heel strike. Left heel strike was determined in real time by detecting when there was a ground reaction force on the left belt and the velocity of the left heel marker first transitioned from positive to negative (Zeni et al., 2008).

### Data analyses

After data were collected, we used custom written MATLAB scripts (v2022a The MathWorks, Natick, MA) to calculate and plot the outcome variables. The primary outcome measure was percent left step length change from baseline (ΔLSL) calculated on each stride (s):

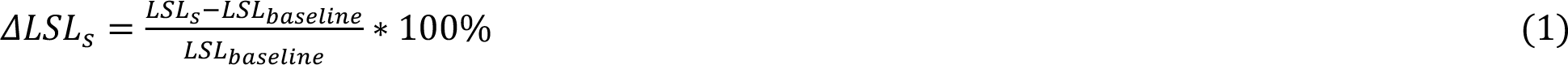

*LSL_baseline_* was calculated as the mean left step length during the final 50 strides of the baseline phase. Key time periods used for analysis during learning (for both experiments) included: early learning, the mean of the first 50 strides after the target reached its plateau (10% change) and late learning, the mean of the last 50 strides of the learning phase.

We tested the hypothesis that the RPE group could learn a new gait pattern by averaging ΔLSL during late learning. Learning in the RPE group would be reflected in a ΔLSL reliably greater than baseline. In addition, to observe how learning emerged in both groups over time, we calculated two secondary outcome measures during early and late learning. Error was calculated as the absolute difference between ΔLSL and the center of the ΔLSL target (10%), and success was calculated as the percentage of left step lengths that fell within the target step length window.

In stable reward environments (in the current study, once the target reached its plateau), exploration should decrease as the values of actions are learned (Daw et al., 2006; Sutton and Barto, 2017). Based on this theory and our assumption that the TE group required little to no exploration, we made two predictions: 1) During early learning, the RPE group would demonstrate greater exploration than the TE group; and 2) The RPE group would reduce their exploration by a greater magnitude from early to late learning compared to the TE group. Our primary measure of exploration, σ_ΔLSL_, was calculated as the standard deviation of ΔLSL, normalized by the ΔLSL standard deviation during the last 50 strides of baseline. Therefore, this measure reflects the increase in motor variability above baseline motor noise. These two predictions served as our test of the hypothesis that the RPE group would demonstrate greater exploration during learning compared to the TE group.

Exploration can also be characterized using a win-stay / lose-shift model. When an individual performs unsuccessful actions, the trial-to-trial changes tend to be more variable than after successful actions (Cashaback et al., 2019; Pekny et al., 2015; Uehara et al., 2019). Therefore, using all of the learning phase data except for the first 50 strides (i.e., before the target started shifting), we performed this trial-to-trial change analysis secondarily to our main exploration analysis. We calculated the difference in ΔLSL values between subsequent strides separately after successful steps and unsuccessful steps. The trial-to-trial change measure was calculated as the standard deviation of these changes, normalized by baseline trial-to-trial change variability (Cashaback et al., 2019; Roth et al., 2023), again to account for increases in this variability relative to baseline. We denote this measure as σ_trial-to-trial_.

For implicit aftereffects and explicit retention, we controlled for any potential differences in learning by calculating both as a percentage of the learned gait pattern for each individual participant on each stride. For experiment 1, we calculated immediate percent implicit aftereffect as the mean of the first 5 strides of the washout phase (immediate washout), and early percent implicit aftereffect as the mean of the next 25 strides (early washout). Therefore, we compared the implicit aftereffects at both timepoints between groups to test the hypothesis that the RPE group would show larger implicit aftereffects than the TE group. For experiment 2, we calculated the percent explicit retention at two different timepoints: the mean of all 25 strides during immediate retention and the mean of the first 25 strides during 24-hour retention. Here, we considered 25 strides so that individuals had time to achieve a consistent level of performance that reflected their explicit motor memory. For experiment 2, it was also necessary to calculate absolute retention error, the absolute difference between ΔLSL at late learning and ΔLSL at immediate and 24-hour retention. This additional measurement of error, or accuracy, was necessary because, unlike for implicit motor memories, we expected that explicit motor memories could either overshoot or undershoot the remembered target location, but the goal was to reproduce the previously learned stepping pattern as accurately as possible. Thus, absolute retention error at both timepoints served as our test for the hypothesis that explicit retention would be more accurate for the RPE group compared to the TE group.

### Statistical analysis

We used Bayesian statistics to make inferences regarding differences within and between groups. This allowed us to estimate the specific probability that our hypotheses were supported or not, given the data we collected as opposed to the classical p value, which provides the probably of obtaining the data, assuming the null hypothesis was true. We assumed data were generated from student’s t distributions with a mean and standard deviation that varied based on both between (group) and within (time) subject factors. We estimated the mean and standard deviation parameters by combining our prior assumptions regarding their values with evidence from the data we collected. We set wide and uninformative priors for each of the model parameters similar to the approached described by Kruschke (2014). We calculated the joint posterior probability distribution with the Python package bayes-toolbox (Kim, 2023) which uses the Pymc library (v4.3; Salvatier et al., 2016) in Python (version 4.3.0). The joint posterior distribution was estimated using Markov Chain Monte Carlo (MCMC) sampling, with 10,000 draws in 4 chains (i.e., 40,000 total samples) and 2,000 tuning samples in each chain. We performed diagnostics for each model, ensuring parameter values were consistent across chains and checking posterior estimates for possible errors (Kruschke, 2014; McElreath, 2016).

We made inferences regarding differences between groups based on posterior contrast distributions: the difference between two posterior distributions of interest. When these contrasts represent a between group difference at a specific timepoint we use the term “group difference”, and when they represent differences across time between groups (analogous to an interaction effect) we use the term “slope difference”. We report the mean difference of each posterior contrast distribution along with the 95% high density interval (HDI), defined as the narrowest span of credible values that contain 95% of the posterior contrast distribution (Kruschke, 2014). We also report the probability of a difference given the data we collected as a percentage of the posterior contrast distribution that lies on one side of 0 (e.g., 85.6% > 0).

### Code Accessibility

The data analysis code for individual participants and figure generation, as well as the full statistical models and code, including detailed information about the priors and diagnostics are available online at the Open Science Framework (https://osf.io/tcwu6/).

## Results

In experiment 1, 15 individuals (9 females; mean age ± 1 SD, 22.8 ± 4.2 years) were randomized into the RPE group and 15 (10 females; 22.5 ± 3.7 years) were randomized into the TE group. In experiment 2, a new sample of individuals was recruited, with 12 (6 females; 23.5 ± 4.6 years) randomized into the RPE group and 12 (7 females; 22.3 ± 4.2 years) randomized into the TE group.

### Experiment 1

#### The RPE group learned the novel locomotor pattern

First, in Figure 2, we display individual learning data. Individual participants displayed in this figure represent their respective groups’ 10^th^, 50^th^ and 90^th^ percentile values for our measure of exploration (σ_ΔLSL_) during early learning. Individuals in the TE group performed consistently with one another across the learning phase, regardless of the percentile (Figure 2, top row), which is not surprising given they received accurate visual feedback of their step length and the target. However, individuals in the RPE group demonstrated a much wider range of idiosyncratic behavior during the learning phase across percentiles (Figure 2, bottom row). The individual representing the 10^th^ percentile performed similar to the individuals in the TE group (Figure 2, bottom left), while the two other individuals show higher variability overall, along with what appears to be different patterns of exploration as learning progressed.

**Figure 2:**
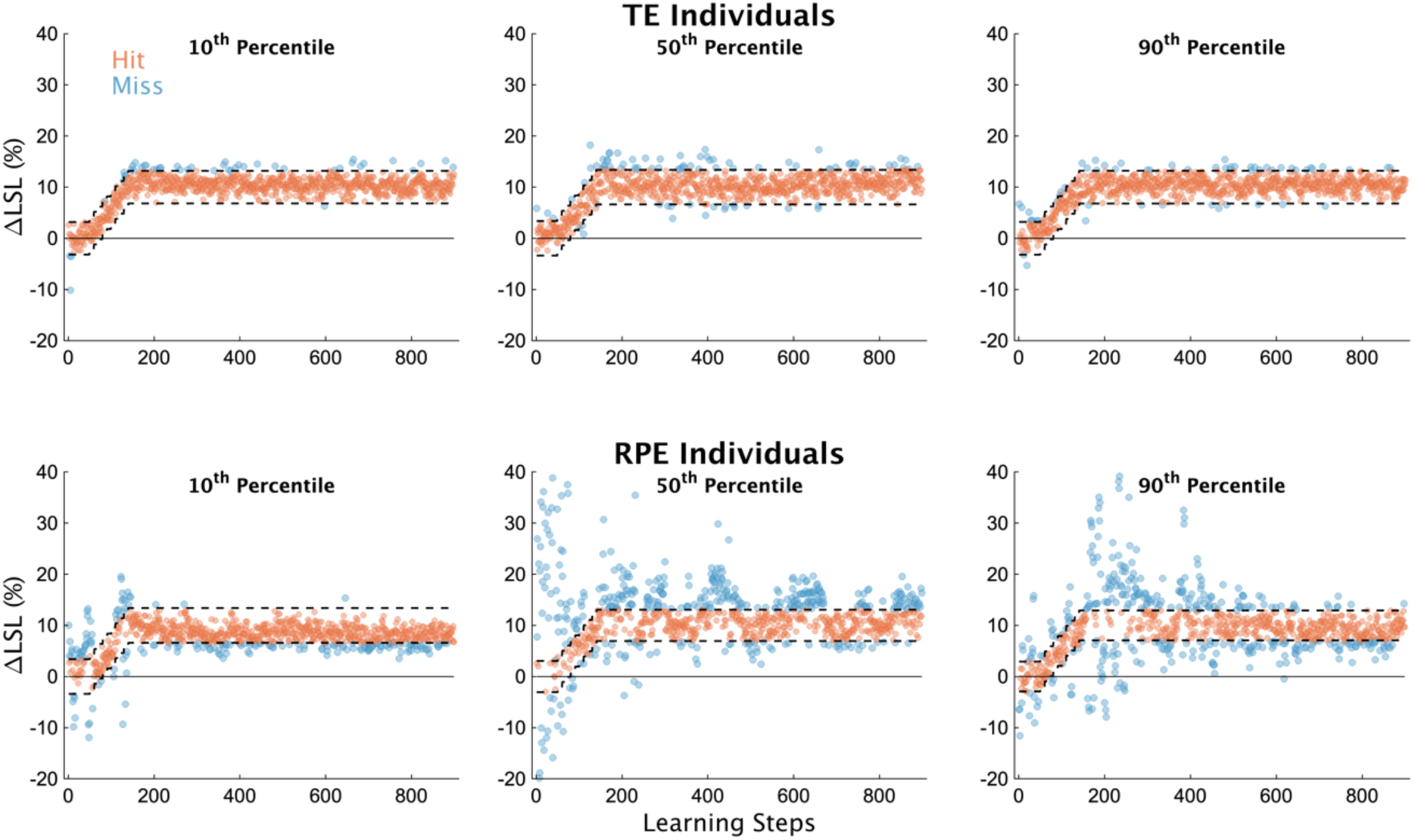
Individual participant data. ΔLSL data for all strides of the learning phase from 6 exemplar subjects; 3 each from the TE and RPE groups. Participants in this figure were selected based on their magnitude of exploration (σ_ΔLSL_ values) during early learning (i.e., the first 50 strides after the target stopped moving). Specifically, we selected individuals who represented the 10^th^, 50^th^ and 90^th^ percentiles for each group separately according to our measure of early exploration. Blue dots represent unsuccessful steps, orange dots represent successful steps. The dashed lines represent the target window, centered on 10% ΔLSL (the width of the window was ±2 cm for all participants).

Despite the idiosyncratic behavior of the RPE group, overall, they learned the new walking pattern by the end of the 900-stride learning phase (Figure 3A). The RPE group’s ΔLSL was much larger than baseline (Figure 3B; RPE late learning, mean [95% HDI] = 9.16% [8.36 9.95], 100% > 0), providing a high degree of certainty that the RPE group learned the new walking pattern. The TE group’s ΔLSL at late learning was slightly greater by comparison (group difference = -0.94% [-1.99 0.02], 96.8% < 0). Additionally, the RPE group’s performance was greatly improved from early to late learning compared to the TE group (Figure 3C and D).

**Figure 3.**
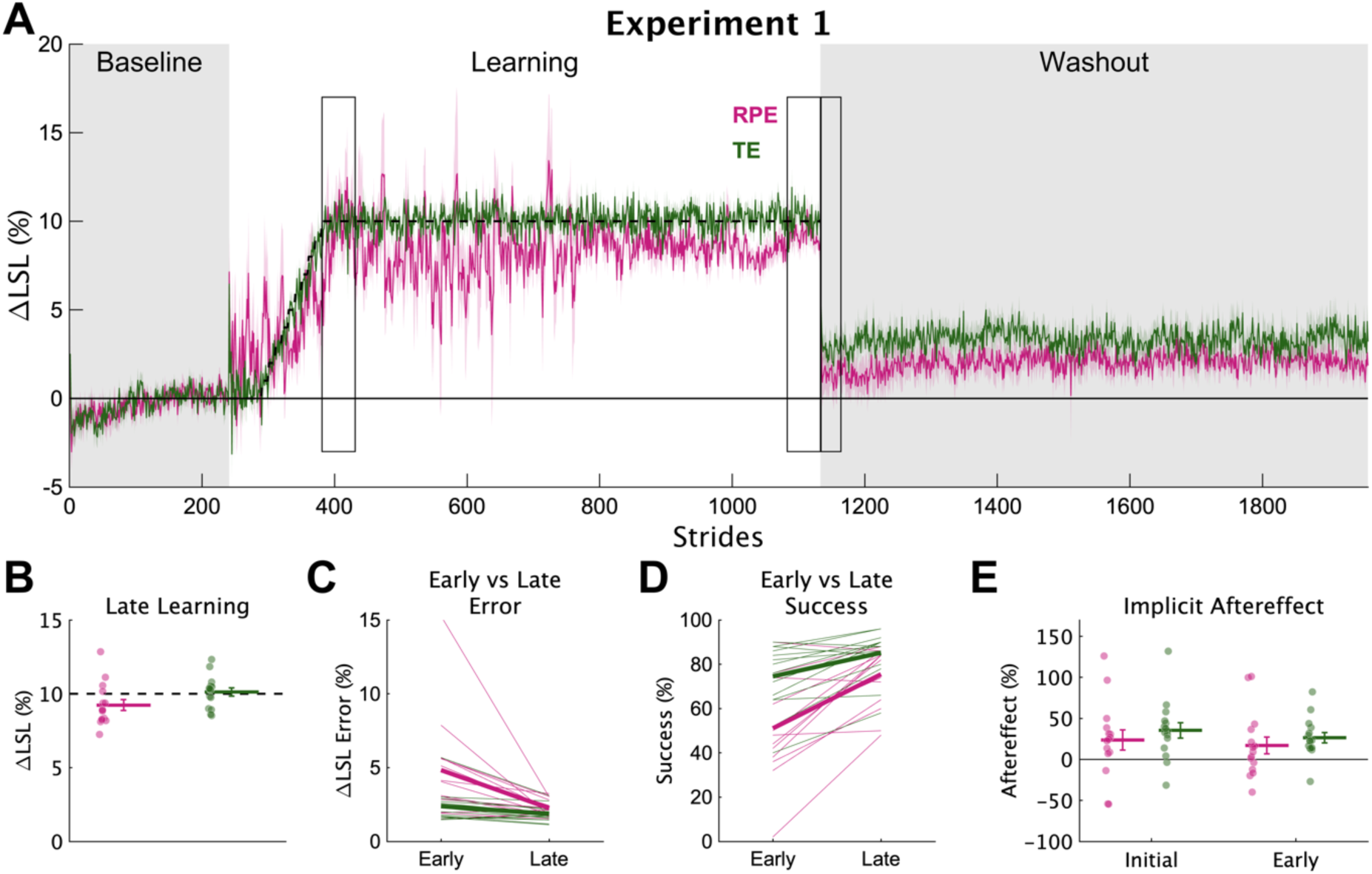
Experiment 1 learning and implicit aftereffects. **(A)** Group average ΔLSL data shown over all strides of experiment 1. Solid colored lines represent group means and shading represents 1 SEM. Gray regions denote the times when no feedback was provided and individuals were asked to “walk normally”. The dashed line represents the center of the target. Rectangular boxes denote the key timepoints of early learning, late learning, and initial and early washout. **(B)** Group averaged ΔLSL at late learning for both groups. Thick horizontal lines represent group means; dots represent individuals; error bars represent ±1 SEM. **(C)** Group average ΔLSL error at early and late learning. Thick lines represent group means; thin lines represent individuals. **(D)** Group average percent success at early and late learning. **(E)** Group average implicit aftereffects. Thick horizontal lines represent group means; dots represent individuals; error bars represent ±1 SEM.

Specifically, compared to the TE group, the RPE group had much larger improvements from early to late learning in terms of error (slope difference = 1.22% [-0.07 2.47], 97.1% > 0) and percent success (slope difference = -12.0% [-23.6 0.8], 97.1% < 0), an effect due to the much larger error in the RPE group in early learning. Together, these findings indicate that the RPE group learned the novel gait pattern, though not as completely, or as quickly, as the TE group.

#### The RPE group explored to learn the novel gait pattern

The results supported the hypothesis that individuals in the RPE group explored to learn the new gait pattern. We found differences in σ_ΔLSL_ between the RPE and TE groups during the learning phase (Figure 4A and B). Specifically, σ_ΔLSL_ was much greater during early learning for individuals in the RPE group compared to individuals in the TE group (group difference = 1.20 [-0.02 2.57], 97.8% > 0), and the σ_ΔLSL_ decreased more from early to late learning for the RPE group compared to the TE group (slope difference = 0.99 [-0.27 2.31], 94.1% > 0). These two results provide strong evidence for the hypothesis that the RPE group explored to learn the new walking pattern.

**Figure 4.**
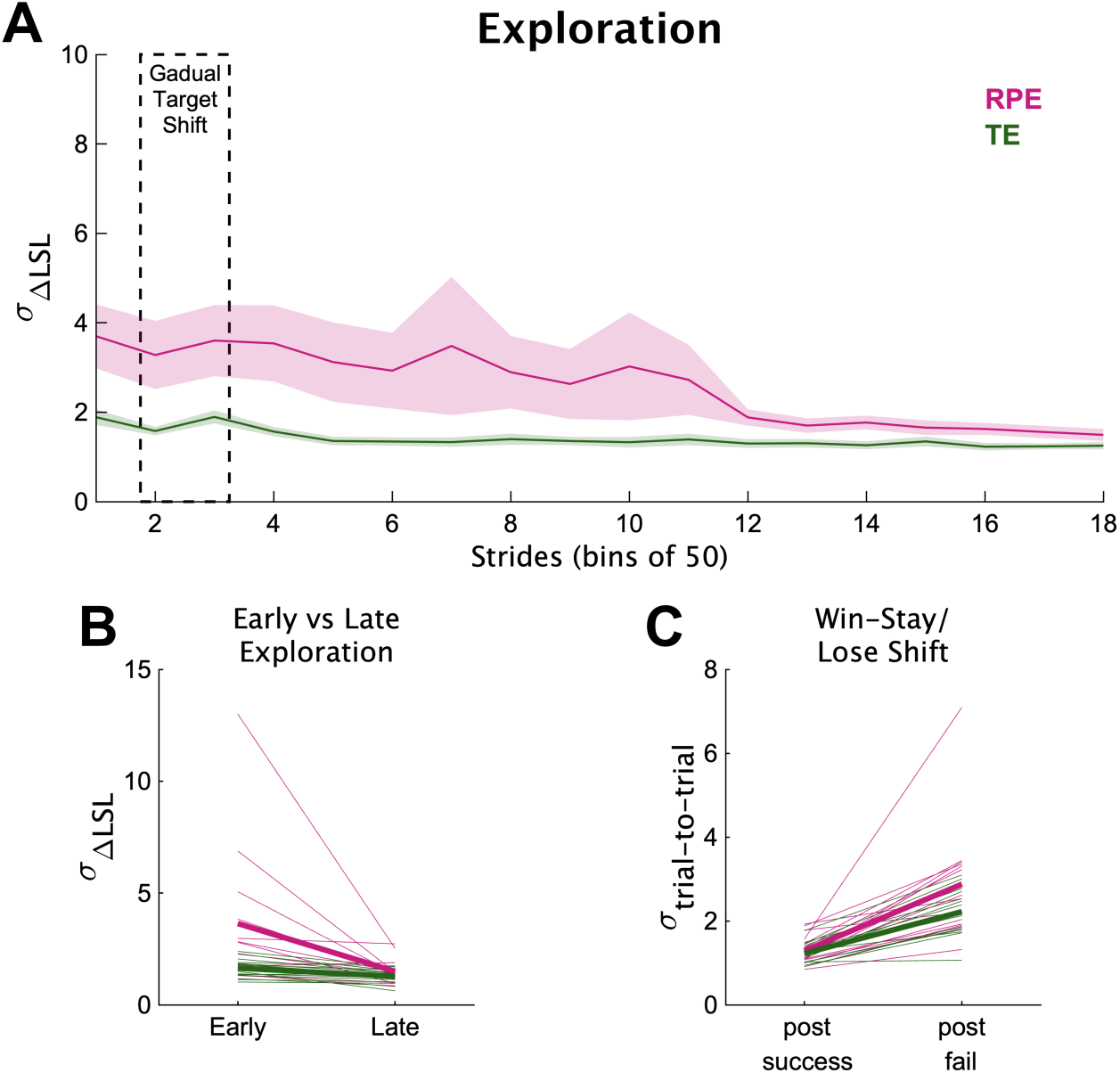
Experiment 1 exploration measurements. **(A)** Exploration across the learning phase, measured as the baseline-normalized standard deviation of ΔLSL. For visualization purposes only, we calculated each individual’s motor variability in 18 bins of 50 strides each across the learning phase, then averaged motor variability among individuals in each group (solid lines) with the shading representing 1 SEM. The dashed rectangle represents the bins when the target was gradually shifting toward 10%. **(B)** Early and late exploration. We calculated σ_ΔLSL_ at early and late learning timepoints. Thick lines represent group means; thin lines represent individuals. **(C)** Motor variability measured as the standard deviation of trial-to-trial changes after successful and unsuccessful steps (σ_trial-to-trial_). Thick lines represent group means; thin lines represent individuals.

In addition to exploration at the two specific timepoints, we also performed a secondary analysis of exploration, measured as the variability of trial-to-trial changes after unsuccessful trials across most of the learning phase (Figure 4C). The RPE group had slightly greater σ_trial-to-trial_ after unsuccessful steps compared to the TE group (group difference = 0.37 [-0.17 1.01], 90.8% > 0). And the difference between σ_trial-to-trial_ for successful vs unsuccessful steps was also slightly larger for the RPE group compared to the TE group (slope difference = 0.31 [-0.25 0.97], 84.6% > 0). Overall, while less reliable than the σ_ΔLSL_ differences (probabilities of ∼91% and 85%), these findings provide additional support for the hypothesis that the RPE group was exploring during locomotion.

#### Reinforcement learning did not increase the magnitude of implicit aftereffects

We probed implicit aftereffects by asking individuals to walk normally during the no-feedback washout phase. Implicit aftereffects were greater for the TE group during initial washout (group difference = -10.7% [-23.8 2.8], 94.9% < 0), and during early washout (group difference = -9.6% [-17.4 -1.9], 98.9% < 0). Therefore, contrary to our hypothesis, learning from reward prediction errors did not increase implicit aftereffects more than learning from target error, rather, the evidence strongly supported the opposite. This finding could reflect a reduced amount of use-dependent learning for the RPE group caused by the increased variability (Wood et al., 2021, 2020).

### Experiment 2

#### Learning from reward prediction errors versus target errors resulted in unique signatures of explicit retention

In experiment 2, we compared explicit retention after learning from reward prediction errors and target errors. This was done with a new sample of individuals who experienced a different post-learning phase but the same baseline and learning phases. We replicated the main findings from experiment 1 (see code for the remainder of the analyses). The RPE group learned (RPE ΔLSL at late learning = 8.74% [7.82 9.66], 100% > 0), and they did so by exploring (group σ_ΔLSL_ difference at early learning = 0.87 [0.07 1.65], 98.6% > 0; σ_ΔLSL_ slope difference = 0.54 [-0.24 1.37], 89.9% > 0). After learning, both groups performed a test of explicit retention where they were instructed to repeat the same walking pattern they performed at the end of learning without feedback, both immediately after learning and 24 hours later (Figure 5A).

**Figure 5.**
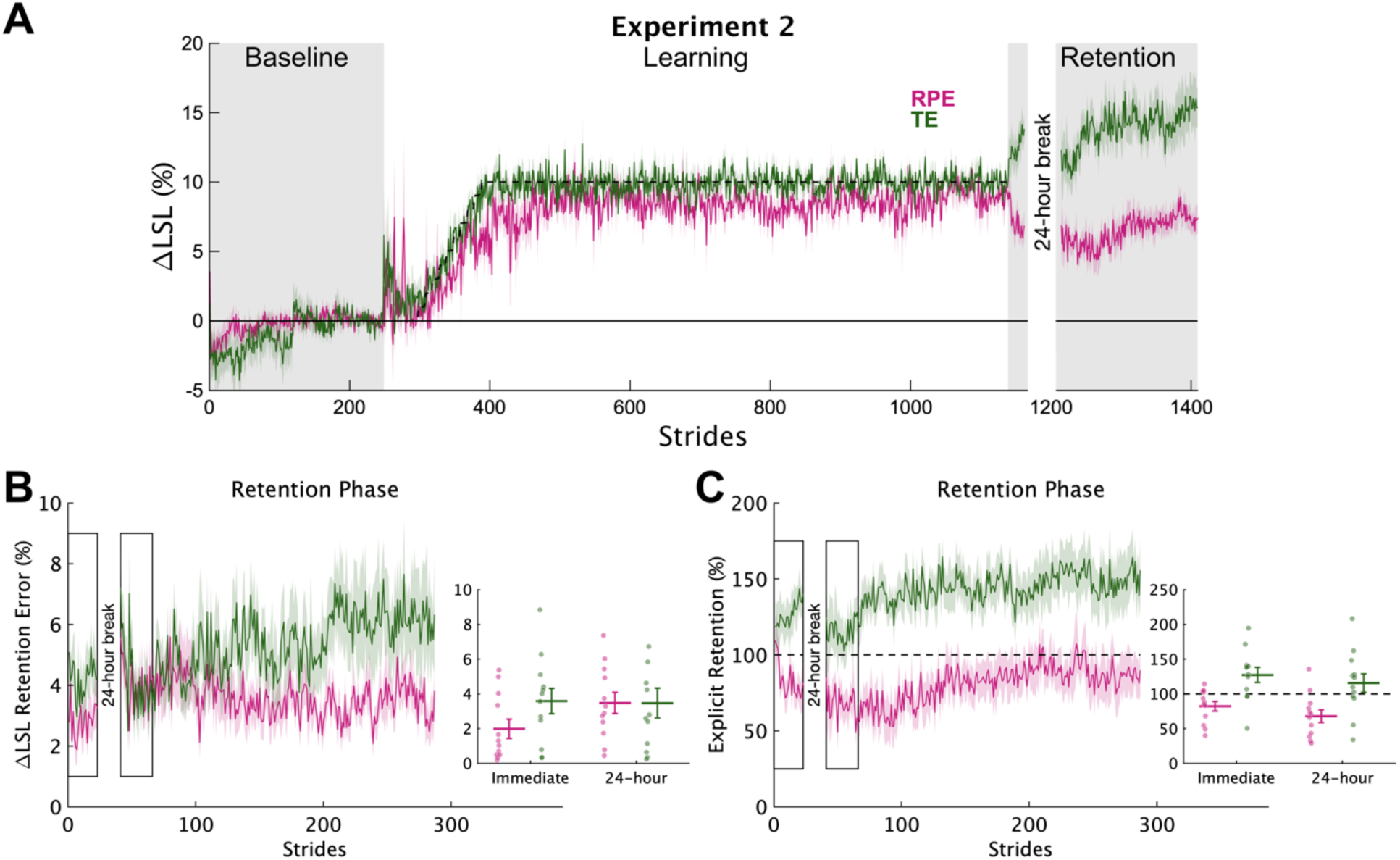
Experiment 2 explicit retention. **(A)** Group average ΔLSL data shown over all strides of experiment 2. Solid colored lines represent group means and shading represents 1 SEM. Gray regions denote the times when no feedback was provided. During retention testing, participants were instructed to “walk like you did at the end of the previous phase”. The dashed line represents the center of the target during the learning phase. **(B)** Group average ΔLSL percent error data for each stride during the immediate and 24-hour retention timepoints (rectangles represent the 25 strides of each epoch). 0 represents perfect retention. The inset shows the group average (horizontal lines) and individual (dots) retention levels for the two timepoints. Error bars represent 1 SEM. **(C)** Group average percent retention data for each stride during the immediate and 24-hour retention timepoints. All shown in the same manner as in **(B)**. The dashed line at 100% represents perfect retention.

First, we assessed the accuracy of the explicit motor memories by comparing the absolute error between the ΔLSL behavior at the end of learning and during the retention phases (Figure 5B). Recall that this is the primary outcome measure of explicit retention because it reflects the true accuracy of retention, or how far each individual’s steps were from the remembered target, regardless of direction. The RPE group demonstrated less absolute retention error at immediate retention compared to the TE group (group difference = -1.36% [-2.94 0.20], 95.8% < 0), but after 24-hours, the RPE group was very similar to the TE group (group difference = -0.13% [-2.01 1.68], 56.5% < 0). Thus, the experiment 2 results revealed that learning a new gait pattern using reward prediction errors produces a unique pattern of explicit retention compared to correcting for target error that is, at least initially, more accurate.

We found large between group differences in the specific walking pattern that was remembered both immediately and 24-hours after learning (Figure 5C). Specifically, percent explicit retention was much larger for the TE group at both immediate (group difference = -44.4% [-63.3 -24.9], 100% < 0) and 24-hour retention (group difference = -45.8% [-71.4 -20.0], 99.8% < 0) compared to the RPE group. Recall that here, “larger” retention does not equate to “better” retention because we asked individuals to walk “exactly” like they did at the end of learning. Rather, this result reflects the fact that the TE group performed a step length that was longer than what they performed at the end of learning (i.e., over-stepping), resulting in percent retention values mostly >100% (9/12 and 8/12 participants for Immediate and 24-hour retention, respectively).

On the other hand, the RPE group performed a step length that was shorter than what they performed at the end of learning (i.e., under-stepping), creating in percent retention values mostly <100% (8/12 and 10/12 participants for Immediate and 24-hour retention, respectively).

#### Explicit retention resulted from differences in declarative knowledge of the successful step length

Why did the RPE and TE groups have such distinct patterns of percent explicit retention? One possibility is that declarative knowledge of the successful step length impacted explicit retention. To investigate this, we used our post-session interviews after the 24-hour retention phase to probe declarative knowledge of successful step lengths. We asked each individual to rate the length of the successful step length compared to their baseline left step into one of three categories: “not longer”, “slightly longer”, or “longer”. The proportions of individuals in each category were as follows: Not longer: TE 0%, RPE 25%; Slightly longer: TE 50%, RPE 50%; Longer: TE 50%, RPE 25%. On average, the TE group rated the successful step length as longer (by about 1 category) than the RPE group (group difference = -1.00 [-2.00 -0.02], 97.7% < 0). Thus, the different feedback and instructions led to differences in declarative knowledge of the successful step length.

## Discussion

This works shows, for the first time, that despite the stability constraints inherent in gait, young healthy individuals can learn a new walking pattern using reward prediction errors. Consistent with the reinforcement learning framework, individuals in the RPE group learned the most valuable action by initially exploring the action space, assigning values to possible step lengths, then as learning progressed, exploiting the learned action/value pairs with more consistent, correct step lengths. We also found that reinforcement learning produced smaller magnitudes of implicit aftereffects, but more accurate explicit retention, than individuals provided with target errors and a specific task strategy, an effect that waned by 24 hours.

### Reinforcement learning during locomotion is accomplished by exploring

Prior studies of locomotor learning have not isolated reward prediction errors by providing only binary reward feedback. Rather, they either added rewards to other teaching signals like target error (Sato et al., 2022) or sensory prediction errors (Bakkum and Marigold, 2022), or used graded reward feedback, increasing the reward as movements became closer to a target (Hasson et al., 2015). While graded reward feedback provides a reward prediction error, having any knowledge about distance or direction from the target adds another learning signal similar to target error. We therefore provided only binary reward feedback in the current study to isolate reinforcement learning as much as possible. The RPE group learned the new walking pattern, a finding consistent with prior studies in reaching (Cashaback et al., 2017; Holland et al., 2018; Izawa and Shadmehr, 2011; Therrien et al., 2016), eye movements (Madelain et al., 2011), and speech production (Parrell, 2021). Unlike a prior study in reaching (Wu et al., 2014), baseline variability was not related to the rate of learning for the RPE groups across both experiments (slope = 0.5 [-2.8 3.8], 63.9% > 0), but our ability to assess the rate of learning could be hindered by the gradual nature of the perturbation in the current study.

To learn the new walking pattern, the RPE group explored the action space. We observed a classic signature of reinforcement learning behavior, the exploration / exploitation trade-off, in the RPE group’s step length variability measurement, σ_ΔLSL_. Interestingly, prior reinforcement-based motor learning studies in human reaching have not observed a reduction in exploratory behavior over time in stable reward environments. We attribute this to the relatively short (360 – 400 trials, Pekny et al., 2015; Uehara et al., 2019), or absent (Cashaback et al., 2017; Izawa and Shadmehr, 2011; Therrien et al., 2016) steady state learning phases in those studies. However, in the current study, steady state learning was 760 strides which allowed the RPE group adequate time to learn the reward landscape well enough to exploit the rewarding step length.

While parsimonious, motor variability is not the only way to characterize exploration in movement-based reinforcement learning experiments. Some methods have characterized exploration as a random walk (Haith and Krakauer, 2014; Roth et al., 2023), but most methods have focused on the magnitude or variability of trial-to-trial of changes after successful vs unsuccessful movements (i.e., win-stay lose shift methods; Pekny et al., 2015; Therrien et al., 2016; Cashaback et al., 2019; Uehara et al., 2019; van Mastrigt et al., 2020, 2021). However, no studies have compared the latter measure of exploration while learning from reward prediction error versus correcting for target errors. While we found that the RPE group had slightly greater trial-to-trial variability after unsuccessful steps compared to the TE group, the effect was relatively small compared to the effect in baseline-normalized motor variability. It is possible other models of exploration, for example, random walk behavior (Roth et al., 2023), may better distinguish between reward prediction error learning and target error correction. Or perhaps these methods of exploration could be task dependent. Future work could elucidate different methods of movement-based exploration similar to the distinct modes of exploration described in decision making tasks (Wilson et al., 2014).

### Reinforcement learning does not increase the magnitude of implicit aftereffects

Reinforcement learning did not increase implicit aftereffects as we expected, and the RPE group actually produced smaller implicit aftereffects compared to the TE group which was not provided with any overt reward. It is possible this difference was due to the increased percent success for the TE group. We performed a Bayesian regression analysis to determine if percent success, measured during the learning phase while the target was at its 10% plateau, could predict immediate implicit aftereffects. We found that the 95% HDIs surrounded a slope value of zero (RPE group slope mean [95% HDI] = 0.32 [-1.56 2.15]; Target group = 0.81 [-0.57 2.27]). Thus, while percent success could have had a positive effect on implicit aftereffects for the Target group, these results are unreliable, a finding that is consistent with our prior work (Wood et al., 2021, 2020).

We suspect the implicit aftereffects in the TE group were caused by use-dependent learning because the feedback was similar to prior studies in our lab and the implicit aftereffect persisted throughout the 900-stride washout phase (Wood et al., 2021, 2020). It is possible that implicit aftereffects in the RPE group were also caused by use-dependent learning as suggested previously (Codol et al., 2018; Holland et al., 2018). If so, the current results contrast somewhat with findings in sensorimotor adaptation (Galea et al., 2015; Huang et al., 2011; Shmuelof et al., 2012; Therrien et al., 2016) and use dependent learning (Floel et al., 2008; Mawase et al., 2017) in which reward boosts implicit aftereffects caused by other learning processes. However, the current results align with work suggesting that increased motor variability attenuates use-dependent learning (Marinovic et al., 2017; Tsay et al., 2022; Verstynen and Sabes, 2011; Wood et al., 2021). Indeed, if we match individuals across groups based on non-normalized, total motor variability (calculated across all strides of the learning phase when the target was plateaued at 10%), the difference in implicit aftereffects disappears (n = 6 per group; initial washout difference = -2.3% [-62.1 61.0], 53.9% < 0; early washout difference = -6.1% [-59.0 47.8], 60.5% < 0). Therefore, we suggest implicit aftereffects in both groups were due to use-dependent learning since, after controlling for motor variability, both groups demonstrated a similar magnitude of implicit aftereffects.

### Reinforcement learning improves the accuracy of explicit retention

Despite less explicit knowledge of the task itself, including the mapping between step length and target position, the RPE group had more accurate immediate explicit retention compared to the TE group. This increased accuracy could have been caused by the reward itself (money and green checkmark) by solidifying the memory of the walking pattern better than simply hitting a target without overt reward. This explanation is similar to Thorndike’s Law of Effect (Thorndike, 1927), and is consistent with studies showing that rewarding successful movements during skill learning and sensorimotor adaptation result in better motor memories than without reward (Abe et al., 2011; Mawase et al., 2017; Shmuelof et al., 2012; Therrien et al., 2016). However, we did not find evidence of a relationship between task success and absolute retention accuracy at immediate retention for either group (RPE = 0.05 [-0.06 0.15]; TE = 0.03 [-0.16 0.20]). Another possibility is that the act of exploring produces a clearer picture of the successful action(s) needed within the workspace, which then leads to more accurate explicit retention. Studies across various learning tasks suggest that greater variability produces worse performance during training but better memories/generalization (Raviv et al., 2022). A recent study in a grid-sailing, key press task suggested that more variable training leads to better knowledge of novel action outcome pair mappings (Velázquez-Vargas et al., 2023). Therefore, it is possible that by exploring the space, individuals in the RPE group developed better knowledge of the action outcome pair.

### Reinforcement learning and correcting for target errors produce distinct explicit motor memories

Interestingly, we found consistent and unique biases in explicit motor memories between groups when assessed as percent explicit retention, a measure that takes the direction of error into account. During both explicit retention phases, the RPE group was biased to step shorter than they did at the end of the learning phase, while the TE group was biased to step longer than they did at the end of the learning phase. Participants’ declarative memories of the successful step length followed a similar pattern (i.e., more individuals in the TE group reported stepping “longer”), suggesting their bias was likely cognitive in nature. While the link between declarative knowledge and different forms of motor learning and memory has been made previously (Haith et al., 2015; Keisler and Shadmehr, 2010; McDougle et al., 2022; Shanks and Johnstone, 1999; Stanley and Krakauer, 2013; Wong et al., 2015), we are not aware of any previous such stark display of distinct biases in declarative motor memories. We speculate that these biases were caused by heuristics, computationally inexpensive mental shortcuts, which not only produce biases in cognitive decisions (Tversky and Kahneman, 1974), but can also explain biases in visual and auditory perception (Branch et al., 2022; Gardner, 2019; Rahnev and Denison, 2018; Raviv et al., 2012).

## Conclusions

Here, we determined that reinforcement learning can drive the acquisition and retention of new gait patterns. Consistent with the reinforcement learning framework, individuals do so by exploring the action space until the most rewarding movement is discovered. Then, the most rewarding action is exploited by repetition. Thus, the reinforcement learning framework can describe performance across a wide range of species and behavior, and bipedal human locomotion is no exception, as our environment is rife with risks (e.g., a trip over a cracked sidewalk) and rewards (e.g., clearing a hurdle on the way to winning a race) that we must learn to pair with specific actions to successfully navigate in the world.

## Acknowledgments

We would like to thank Saunders Penn for his help with data collections and making Figure 1A and Maci Keaton for her help with data collections.

## References

1. Abe M, Schambra H, Wassermann EM, Luckenbaugh D, Schweighofer N, Cohen LG (2011) Reward Improves Long-Term Retention of a Motor Memory through Induction of Offline Memory Gains. Current Biology 21:557–562.

2. Bakkum A, Marigold DS (2022) Learning from the Physical Consequences of Our Actions Improves Motor Memory. eNeuro 9.

3. Bao S, Lei Y (2022) Memory decay and generalization following distinct motor learning mechanisms. Journal of Neurophysiology 128:1534–1545.

4. Brach JS, Berlin JE, VanSwearingen JM, Newman AB, Studenski SA (2005) Too much or too little step width variability is associated with a fall history in older persons who walk at or near normal gait speed. J Neuroeng Rehabil 2:21.

5. Branch F, Park E, Hegdé J (2022) Heuristic Vetoing: Top-Down Influences of the Anchoring-and-Adjustment Heuristic Can Override the Bottom-Up Information in Visual Images. Front Neurosci 16:745269.

6. Cashaback JGA, Lao CK, Palidis DJ, Coltman SK, McGregor HR, Gribble PL (2019) The gradient of the reinforcement landscape influences sensorimotor learning. PLoS Comput Biol 15:e1006839.

7. Cashaback JGA, McGregor HR, Mohatarem A, Gribble PL (2017) Dissociating error-based and reinforcement-based loss functions during sensorimotor learning. PLOS Computational Biology 13:e1005623.

8. Codol O, Holland PJ, Galea JM (2018) The relationship between reinforcement and explicit control during visuomotor adaptation. Scientific Reports 8.

9. Daw ND, O’Doherty JP, Dayan P, Seymour B, Dolan RJ (2006) Cortical substrates for exploratory decisions in humans. Nature 441:876–879.

10. Dhawale AK, Smith MA, Ölveczky BP (2017) The Role of Variability in Motor Learning. Annu Rev Neurosci 40:479–498.

11. Floel A, Garraux G, Xu B, Breitenstein C, Knecht S, Herscovitch P, Cohen LG (2008) Levodopa increases memory encoding and dopamine release in the striatum in the elderly. Neurobiol Aging 29:267–279.

12. French MA, Morton SM, Charalambous CC, Reisman DS (2018) A locomotor learning paradigm using distorted visual feedback elicits strategic learning. J Neurophysiol 120:1923–1931.

13. French MA, Morton SM, Reisman DS (2021) Use of explicit processes during a visually guided locomotor learning task predicts 24-h retention after stroke. Journal of Neurophysiology 125:211–222.

14. Galea JM, Mallia E, Rothwell J, Diedrichsen J (2015) The dissociable effects of punishment and reward on motor learning. Nature Neuroscience 18:597–602.

15. Gardner JL (2019) Optimality and heuristics in perceptual neuroscience. Nat Neurosci 22:514– 523.

16. Haith A, Krakauer J (2014) Motor Learning by Sequential Sampling of Actions, Presented at the Translational and Computational Motor Control. p2. Washington D.C.

17. Haith AM, Huberdeau DM, Krakauer JW (2015) The Influence of Movement Preparation Time on the Expression of Visuomotor Learning and Savings. Journal of Neuroscience 35:5109–5117.

18. Hasson CJ, Manczurowsky J, Yen S-C (2015) A reinforcement learning approach to gait training improves retention. Frontiers in Human Neuroscience 9.

19. Hausdorff JM, Edelberg HK, Mitchell SL, Goldberger AL, Wei JY (1997) Increased gait unsteadiness in community-dwelling elderly fallers. Archives of Physical Medicine and Rehabilitation 78:278–283.

20. Hausdorff JM, Rios DA, Edelberg HK (2001) Gait variability and fall risk in community-living older adults: A 1-year prospective study. Archives of Physical Medicine and Rehabilitation 82:1050–1056.

21. Holland P, Codol O, Galea JM (2018) Contribution of explicit processes to reinforcement-based motor learning. Journal of Neurophysiology 119:2241–2255.

22. Huang VS, Haith A, Mazzoni P, Krakauer JW (2011) Rethinking motor learning and savings in adaptation paradigms: model-free memory for successful actions combines with internal models. Neuron 70:787–801.

23. Izawa J, Shadmehr R (2011) Learning from sensory and reward prediction errors during motor adaptation. PLoS Comput Biol 7:e1002012.

24. Keisler A, Shadmehr R (2010) A Shared Resource between Declarative Memory and Motor Memory. J Neurosci 30:14817–14823.

25. Kim HE (2023) bayes-toolbox.

26. Krakauer JW, Hadjiosif AM, Xu J, Wong AL, Haith AM (2019) Motor Learning In: Comprehensive Physiology, pp 613–663. American Cancer Society.

27. Kruschke J (2014) Doing Bayesian Data Analysis: A Tutorial with R, JAGS, and Stan. Academic Press.

28. Madelain L, Paeye C, Wallman J (2011) Modification of saccadic gain by reinforcement. Journal of Neurophysiology 106:219–232.

29. Maki BE (1997) Gait changes in older adults: predictors of falls or indicators of fear. J Am Geriatr Soc 45:313–320.

30. Manley H, Dayan P, Diedrichsen J (2014) When Money Is Not Enough: Awareness, Success, and Variability in Motor Learning. PLoS ONE 9:e86580.

31. Marinovic W, Poh E, de Rugy A, Carroll TJ (2017) Action history influences subsequent movement via two distinct processes. eLife 6:e26713.

32. Mawase F, Uehara S, Bastian AJ, Celnik P (2017) Motor Learning Enhances Use-Dependent Plasticity. The Journal of Neuroscience 37:2673–2685.

33. McAndrew Young PM, Dingwell JB (2012) Voluntarily changing step length or step width affects dynamic stability of human walking. Gait & Posture 35:472–477.

34. McDougle SD, Wilterson SA, Turk-Browne NB, Taylor JA (2022) Revisiting the Role of the Medial Temporal Lobe in Motor Learning. Journal of Cognitive Neuroscience 34:532– 549.

35. McElreath R (2016) Statistical Rethinking: A Bayesian Course with Examples in R and Stan. CRC Press.

36. Parrell B (2021) A Potential Role for Reinforcement Learning in Speech Production. Journal of Cognitive Neuroscience 1–17.

37. Pekny SE, Izawa J, Shadmehr R (2015) Reward-Dependent Modulation of Movement Variability. Journal of Neuroscience 35:4015–4024.

38. Rahnev D, Denison RN (2018) Suboptimality in perceptual decision making. Behavioral and Brain Sciences 41:e223.

39. Raviv L, Lupyan G, Green SC (2022) How variability shapes learning and generalization. Trends in Cognitive Sciences 26:462–483.

40. Raviv O, Ahissar M, Loewenstein Y (2012) How Recent History Affects Perception: The Normative Approach and Its Heuristic Approximation. PLoS Comput Biol 8:e1002731.

41. Reisman DS, Block HJ, Bastian AJ (2005) Interlimb coordination during locomotion: what can be adapted and stored? J Neurophysiol 94:2403–2415.

42. Rescorla R, Wagner A (1972) A theory of Pavlovian conditioning: The effectiveness of reinforcement and non-reinforcement. Classical Conditioning: Current Research and Theory.

43. Roemmich RT, Long AW, Bastian AJ (2016) Seeing the Errors You Feel Enhances Locomotor Performance but Not Learning. Current Biology 26:2707–2716.

44. Roth AM, Calalo JA, Lokesh R, Sullivan SR, Grill S, Jeka JJ, van der Kooij K, Carter MJ, Cashaback JGA (2023) Reinforcement-Based Processes Actively Regulate Motor Exploration Along Redundant Solution Manifolds (preprint). Neuroscience.

45. Salvatier J, Wiecki TV, Fonnesbeck C (2016) Probabilistic programming in Python using PyMC3. PeerJ Comput Sci 2:e55.

46. Sato S, Cui A, Choi JT (2022) Visuomotor errors drive step length and step time adaptation during ‘virtual’ split-belt walking: the effects of reinforcement feedback. Exp Brain Res 240:511–523.

47. Schmidt RA, Lee TD (2005) Motor control and learning: A behavioral emphasis, 4th ed, Motor control and learning: A behavioral emphasis, 4th ed. Champaign, IL, US: Human Kinetics.

48. Shanks DR, Johnstone T (1999) Evaluating the relationship between explicit and implicit knowledge in a sequential reaction time task. Journal of Experimental Psychology: Learning, Memory, and Cognition 25:1435–1451.

49. Shmuelof L, Huang VS, Haith AM, Delnicki RJ, Mazzoni P, Krakauer JW (2012) Overcoming motor “forgetting” through reinforcement of learned actions. J Neurosci 32:14617– 14621a.

50. Stanley J, Krakauer JW (2013) Motor skill depends on knowledge of facts. Front Hum Neurosci 7.

51. Sutton R, Barto A (2017) Reinforcement Learning: An Introduction, 2nd ed. Cambridge, Massachusetts: MIT Press.

52. Therrien AS, Wolpert DM, Bastian AJ (2018) Increasing Motor Noise Impairs Reinforcement Learning in Healthy Individuals. eneuro 5:ENEURO.0050-18.2018.

53. Therrien AS, Wolpert DM, Bastian AJ (2016) Effective reinforcement learning following cerebellar damage requires a balance between exploration and motor noise. Brain 139:101–114.

54. Thorndike EL (1927) The Law of Effect. The American Journal of Psychology 39:212–222.

55. Tsay JS, Kim HE, Saxena A, Parvin DE, Verstynen T, Ivry RB (2022) Dissociable use-dependent processes for volitional goal-directed reaching. Proc R Soc B 289:20220415.

56. Tversky A, Kahneman D (1974) Judgment under Uncertainty: Heuristics and Biases. Science 185:1124–1131.

57. Uehara S, Mawase F, Therrien AS, Cherry-Allen KM, Celnik P (2019) Interactions between motor exploration and reinforcement learning. Journal of Neurophysiology 122:797–808.

58. van Mastrigt NM, Smeets JBJ, van der Kooij K (2020) Quantifying exploration in reward-based motor learning. PLoS ONE 15:e0226789.

59. van Mastrigt NM, van der Kooij K, Smeets JBJ (2021) Pitfalls in quantifying exploration in reward-based motor learning and how to avoid them. Biol Cybern 115:365–382.

60. Velázquez-Vargas CA, Daw ND, Taylor JA (2023) Learning generalizable visuomotor mappings for de novo skills.

61. Verstynen T, Sabes PN (2011) How each movement changes the next: an experimental and theoretical study of fast adaptive priors in reaching. J Neurosci 31:10050–10059.

62. Wilson RC, Geana A, White JM, Ludvig EA, Cohen JD (2014) Humans use directed and random exploration to solve the explore–exploit dilemma. Journal of Experimental Psychology: General 143:2074–2081.

63. Winstein CJ, Pohl PS, Lewthwaite R (1994) Effects of physical guidance and knowledge of results on motor learning: support for the guidance hypothesis. Res Q Exerc Sport 65:316–323.

64. Winstein CJ, Schmidt RA (1990) Reduced frequency of knowledge of results enhances motor skill learning. Journal of Experimental Psychology: Learning, Memory, and Cognition 16:677–691.

65. Wong AL, Lindquist MA, Haith AM, Krakauer JW (2015) Explicit knowledge enhances motor vigor and performance: motivation versus practice in sequence tasks. Journal of Neurophysiology 114:219–232.

66. Wood JM, Kim HE, French MA, Reisman DS, Morton SM (2020) Use-dependent plasticity explains aftereffects in visually guided locomotor learning of a novel step length asymmetry. Journal of Neurophysiology 124:32–39.

67. Wood JM, Morton SM, Kim HE (2021) The Consistency of Prior Movements Shapes Locomotor Use-Dependent Learning. eNeuro 8.

68. Wu HG, Miyamoto YR, Castro LNG, Ölveczky BP, Smith MA (2014) Temporal structure of motor variability is dynamically regulated and predicts motor learning ability. Nat Neurosci 17:312–321.

69. Zeni JA, Richards JG, Higginson JS (2008) Two simple methods for determining gait events during treadmill and overground walking using kinematic data. Gait & Posture 27:710– 714.

